# Proteomic Investigation of the Signal Transduction Pathways Controlling Colistin Resistance in *Klebsiella pneumoniae*

**DOI:** 10.1101/2020.05.05.078428

**Authors:** Ching Hei Phoebe Cheung, Punyawee Dulyayangkul, Kate J. Heesom, Matthew B. Avison

**Affiliations:** School of Cellular & Molecular Medicine, University of Bristol, Bristol. UK; University of Bristol Proteomics Facility, Bristol, UK

## Abstract

Colistin resistance in *Klebsiella pneumoniae* is predominantly caused by mutations that increase expression of the *arn* (also known as *pbg* or *pmrF*) operon. Expression is activated by the PhoPQ and PmrAB two-component systems. Constitutive PhoPQ activation occurs directly by mutation or following loss of MgrB. PhoPQ may also cross-activate PmrAB via the linker protein PmrD. Using proteomics, we show that MgrB loss causes a wider proteomic effect than direct PhoPQ activation, suggesting additional targets for MgrB. Different *mgrB* mutations cause different amounts of Arn protein production, which correlated with colistin MIC. Disruption of *phoP* in an *mgrB* mutant had a reciprocal effect to direct activation of PhoQ in a wild-type background, but the regulated proteins showed almost total overlap. Disruption of *pmrD* or *pmrA* slightly reduced Arn protein production in an *mgrB* mutant, but production was still high enough to confer colistin resistance; disruption of *phoP* conferred wild-type Arn production and colistin MIC. Activation of PhoPQ directly, or through *mgrB* mutation did not significantly activate PmrAB or PmrC production but direct activation of PmrAB by mutation did, and also activated Arn production and conferred colistin resistance. There was little overlap between the PmrAB and PhoPQ regulons. We conclude that under the conditions used for colistin susceptibility testing, PhoPQ-PmrD-PmrAB cross-regulation is not significant and that independent activation of PhoPQ or PmrAB is the main reason that Arn protein production increases above the threshold required for colistin resistance.

## Introduction

Colistin is increasingly used to treat infections caused by extensively drug resistant Gram-negative bacteria (1). Colistin resistance in carbapenem-resistant *Klebsiella pneumoniae*, which was first reported in 2010 (2–4) is, therefore, a critically important problem. It can be caused by mobile *mcr* genes but by far the most common causes are chromosomal mutations (5,6). For example, loss-of-function mutations in *mgrB* regularly emerge following colistin therapy in the clinic, and when selecting resistant mutants in the laboratory (7–10). Loss of MgrB causes activation of *arn* operon expression (7). This operon, also referred to as the *pbg* or *pmrF* operon (5) encodes a series of enzymes forming a pathway that modifies lipid A in lipopolysaccharide by adding 4‐amino‐4‐deoxy‐L‐arabinose. This modification has been seen in colistin resistant *K. pneumoniae* mutants in many studies (e.g. 11–13). Its effect is to reduce cell surface negative charge, reducing affinity for positively charged colistin, raising its MIC (5).

Activation of *arn* operon transcription in *K. pneumoniae* involves two upstream promoters, each targeted by a two-component system response regulator. PhoP targets promoter 1 in response to low magnesium concentrations and PmrA targets promoter 2 in response to high iron concentrations and low pH (14). The dramatic impact that cations and pH have on *arn* promoter activity explains the wide range of medium-dependent colistin MICs observed in the laboratory (15,16). According to a joint CLSI and EUCAST report, colistin susceptibility testing needs to be tightly standardised, therefore, and the gold standard is broth microdilution using colistin sulphate, cation adjusted Muller Hinton broth and with no additives (17)

Additional complexity arises in control of *arn* operon transcription in *K. pneumoniae* because the response regulator PhoP can also activate transcription of *pmrD* and PmrD binds PmrA and enhances its activation (14,18). Hence, when low magnesium and high iron occur at the same time, the PmrA-targeted *arn* operon promoter 2 is more strongly activated than it is in the presence of high magnesium and high iron (14).

The cognate sensor kinases activating PhoP and PmrA are PhoQ and PmrB, respectively (14). It is generally accepted that loss-of-function mutations in *mgrB* activate *arn* operon transcription in *K. pneumoniae* because MgrB is a direct negative regulator of PhoQ sensor kinase activity (5). In fact, this is experimentally confirmed only in *Salmonella* spp. (19) but loss of MgrB does constitutively activate PhoPQ in *K. pneumoniae*, leading to constitutively enhanced transcription from *arn* operon promoter 1 (14). Specific mutations in PhoPQ also activate *arn* operon transcription and confer colistin resistance or heteroresistance in *K. pneumoniae*, e.g. PhoQ substitutions Asp434Asn (20) or Ala21Ser (21) or PhoP substitution Asp191Tyr (22).

Mutations in PmrAB, constitutively activating *arn* operon transcription from promoter 2 have also been found to cause colistin resistance in *K. pneumoniae*. For example, Leu82Arg (23) or Thr157Pro (24) in PmrB, but activation of PmrAB also increases expression of *pmrC*. This encodes an enzyme that modifies lipopolysaccharide by decorating it with phosphoethanolamine, which contributes to colistin resistance in *Salmonella* spp. because it is another way to reduce negative charge on the cell surface (25).

A third two-component regulatory system known to be involved in colistin resistance in *K. pneumoniae* is CrrAB. Activatory mutations in the sensor kinase CrrB have been identified in colistin resistant clinical isolates (20,26,27). Based on transcriptomic analysis these mutants have increased *arn* operon and *pmrC* transcription (20) and in this they closely resemble PmrAB activatory mutants and indeed *pmrAB* is essential for the activation of *arn* operon and *pmrC* transcription in CrrAB activatory mutants, suggesting direct linkage between these two-component systems (26). One additional effect of CrrAB activation is upregulation of *crrC* transcription (20) and *crrC* is also essential for activation of *arn* operon and *pmrC* transcription in a CrrAB activatory mutant (26). This suggests that CrrC forms the link between activated CrrAB and activation of PmrAB, leading to colistin resistance, but the mechanism by which this link operates is not yet known.

The complexity associated with acquisition of colistin resistance in *K. pneumoniae*, means that clinical cases involve a wide range of mutations. In a recent large clinical survey, *mgrB* loss-of-function was the most common mechanism, but in many cases, additional mutations in two-component system genes was seen, and in cases where multiple mutations were seen, a sequential increase in colistin MIC was observed (28)

The aim of the work reported here was to use LC-MS/MS shotgun proteomics and targeted mutagenesis to investigate the importance of the MgrB-PhoPQ-PmrD-PmrAB signal transduction pathway to modulate Arn protein and PmrC production, and assess the protein abundance thresholds required for colistin resistance in *K. pneumoniae* stimulated by mutations affecting PhoPQ and/or PmrAB activity.

## Results and Discussion

### The direct role of PhoPQ and the indirect role of signal transduction from PhoPQ through PmrD to PmrAB in Arn protein production and colistin resistance caused by mgrB mutation

A collection of six *K. pneumoniae* spontaneous single-step mutants were selected in the laboratory using Muller-Hinton agar containing 32 μg.mL^−1^ of colistin, and in each case, PCR sequencing confirmed mutation in *mgrB* or upstream. Three different mutations were seen, causing the following changes: Gln29STOP in MgrB (found in four colistin resistant mutants here, represented by mutant P21); an A to G transition at −31 relative to the *mgrB* start codon, weakening a putative second −10 promoter sequence (**Figure 1**) (represented by mutant P22); a deletion comprising the region between −19 relative to the start codon, to remove the first 41 amino acids of MgrB (represented by mutant P23).

**Figure 1.**
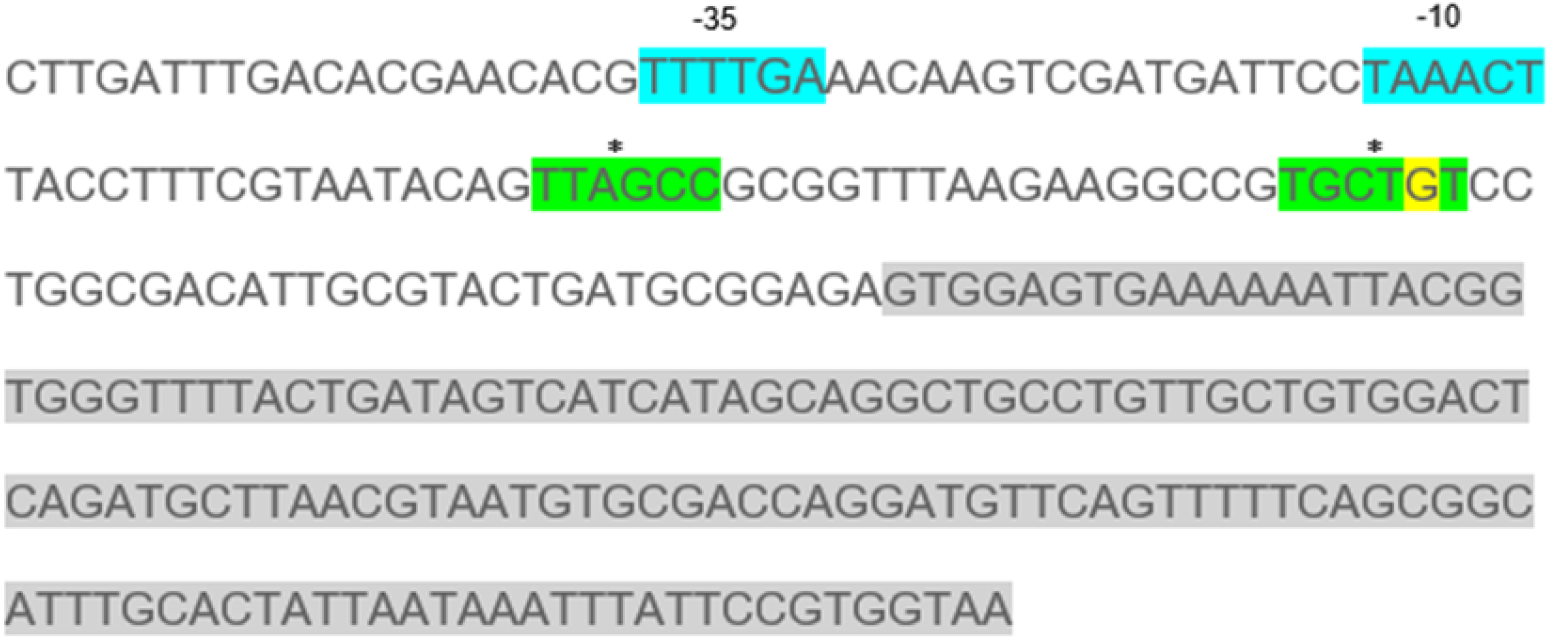
Sequence at the 5’ end of *mgrB* and upstream in colistin resistant mutant P22. The two promoters for *mgrB* are highlighted in blue and green. The mutation in P22 alters the wild-type sequence at the position highlighted in yellow from A to G in the putative −10 box of the second promoter, which is marked with two stars. The first promoter sequence, defined as −35 and −10 was defined in (10).

Colistin MICs against these three representative mutants were determined and, in all cases, colistin resistance was confirmed (**Table 1**). Envelope proteomics identified 45 proteins significantly differentially regulated in all three *mgrB* mutants relative to the parent strain, Ecl8 (33 up- and 12 down-regulated; **Table 2**). These included the ArnABCDT operon proteins known to be responsible for modification of lipopolysaccharide by the addition of 4-amino-4-deoxy-L-arabinose (5). **Figure 2**shows that Arn protein production is highest in the mutant P23, where MgrB is entirely lost. The *mgrB* nonsense mutant truncated at 29 amino acids, P22 and the *mgrB* promoter mutant P21 both have significantly lower Arn protein production than in P23, implying some residual repressive activity of MgrB in both cases. Overall, P22 had the lowest Arn protein production of the three (**Figure 2**). Nonetheless all three mutants are colistin resistant though, as expected based on Arn protein production levels, the highest MIC is against P23 and the lowest against P22 (**Table 1**). This leads to the conclusion that once Arn protein production increases above a certain threshold, colistin resistance is conferred, but that as protein production increases further – P23>P21>P22 – MIC also increases.

**Table 1:** MICs of colistin against clinical isolates and mutant derivatives. Values reported are the modes of three repetitions. Shading indicates resistance according to susceptibility breakpoints set by the CLSI (35).

| Strain/ mutant | Colistin MIC ($\mu\text{g.mL}^{-1}$) |
| --- | --- |
| Ecl8 | 1 |
| P21 (*mgrB*) | 64 |
| P22 (*mgrB*) | 32 |
| P23 (*mgrB*) | 128 |
| P23 *phoP* | 2 |
| P23 *pmrD* | 64 |
| P23 *pmrA* | 64 |
| KP47 | 2 |
| KP47 PhoQ* | 64 |
| KP47 PmrB* | 32 |

**Table 2:** Significant changes in envelope protein abundance seen in *K. pneumoniae mgrB* mutant P21 versus parent strain Ecl8. Strains were grown in CAMHB and raw abundance data for each protein in a sample are presented normalised using the average abundance the 50 most abundant proteins in that sample. Data for three biological replicates of parent (Ecl8) and *mgrB* mutant (P21). Proteins listed are se significantly differently up- (fold change >1) or down-regulated (fold change <1) in P21, and in *mgrB* mutants P22 and P23 versus their parent Ecl8, based on t-test. Shading indicates proteins also upregulated in the PmrB* (activatory) mutant of *K. pneumoniae* clinical isolate 47 relative to KP47, see text. Stars indicate proteins encoded by transcripts upregulated in a *K. pneumoniae* CrrB activatory mutant (20).

| Accession | Description | Ecl8 1 | Ecl8 2 | Ecl8 3 | P21 1 | P21 2 | P21 3 | Fold Change | t-test |
| --- | --- | --- | --- | --- | --- | --- | --- | --- | --- |
| A6T5Y8 | Mg2+ transport ATPase | 0.01 | 0.00 | 0.00 | 0.02 | 0.04 | 0.03 | 7.63 | 0.01 |
| A6T7F8 | Thymidylate kinase | 0.01 | 0.00 | 0.00 | 0.01 | 0.02 | 0.01 | 4.29 | 0.02 |
| A6TBQ4 | Lipid A 1-diphosphate synthase | 0.00 | 0.00 | 0.00 | 0.07 | 0.09 | 0.08 | >100 | 0.00 |
| A6THT5 | Putative porin | 0.01 | 0.02 | 0.00 | 0.16 | 0.16 | 0.08 | 13.77 | 0.01 |
| A6T4W9 | Bifunctional uridylyltransferase/<br>uridylyl-removing enzyme | 0.00 | 0.00 | 0.00 | 0.01 | 0.01 | 0.00 | 2.76 | 0.04 |
| A6T5U9 | Putative periplasmic binding protein | 0.00 | 0.00 | 0.00 | 0.01 | 0.02 | 0.01 | >100 | <0.005 |
| A6THH1 | MgtA | 0.00 | 0.00 | 0.00 | 0.01 | 0.03 | 0.02 | >100 | <0.005 |
| A6T6X6 | MacA | 0.00 | 0.00 | 0.00 | 0.01 | 0.02 | 0.02 | 11.68 | 0.01 |
| A6T7D1 | Hypothetical Protein | 0.01 | 0.00 | 0.01 | 0.01 | 0.01 | 0.01 | 1.83 | 0.03 |
| A6T7J8 | PhoP | 0.01 | 0.01 | 0.01 | 0.05 | 0.06 | 0.05 | 5.41 | <0.005 |
| A6T9Y9 | SlyB | 0.37 | 0.38 | 0.17 | 0.81 | 1.04 | 0.46 | 2.52 | 0.03 |
| A6TBT1 | AbpE | 0.00 | 0.01 | 0.00 | 0.01 | 0.03 | 0.01 | 5.61 | 0.02 |
| A6TCT2 | LpxO | 0.01 | 0.01 | 0.01 | 0.03 | 0.04 | 0.02 | 4.77 | 0.01 |
| A6TF96 | ArnT | 0.00 | 0.00 | 0.00 | 0.02 | 0.01 | 0.01 | >100 | <0.005 |
| A6TF97 | ArnD | 0.00 | 0.00 | 0.00 | 0.03 | 0.04 | 0.02 | >100 | <0.005 |
| A6TF98 | ArnA | 0.01 | 0.00 | 0.00 | 0.09 | 0.11 | 0.15 | 20.27 | <0.005 |
| A6TF99 | ArnC | 0.00 | 0.00 | 0.00 | 0.05 | 0.08 | 0.05 | >100 | <0.005 |
| A6TFA0 | ArnB | 0.00 | 0.00 | 0.00 | 0.05 | 0.00 | 0.04 | >100 | 0.04 |
| A6T7H3 | Putative enzyme | 0.01 | 0.01 | 0.01 | 0.01 | 0.01 | 0.01 | 1.67 | 0.04 |
| A6TEP0 | Cell division protein ZapE | 0.02 | 0.01 | 0.01 | 0.05 | 0.04 | 0.02 | 2.36 | 0.03 |
| A6T6G3 | Tol-Pal system protein TolQ | 0.04 | 0.04 | 0.03 | 0.06 | 0.08 | 0.05 | 1.73 | 0.04 |
| A6TDS9 | Component of the MscS mechanosensitive channel | 0.06 | 0.03 | 0.04 | 0.06 | 0.09 | 0.07 | 1.65 | 0.04 |
| A6T4X0 | Methionine aminopeptidase | 0.04 | 0.04 | 0.05 | 0.06 | 0.05 | 0.05 | 1.29 | 0.01 |
| A6T5I4 | Peptidylprolyl isomerase | 0.13 | 0.07 | 0.09 | 0.20 | 0.18 | 0.21 | 2.00 | <0.005 |
| A6T642 | 2,3-dihydro-2,3-dihydroxybenzoate synthetase, isochroismatase | 0.00 | 0.00 | 0.01 | 0.01 | 0.01 | 0.01 | 2.17 | 0.03 |
| A6T7K1 | Adenylosuccinate lyase | 0.16 | 0.09 | 0.12 | 0.16 | 0.22 | 0.17 | 1.47 | 0.05 |
| A6T7R3 | Putative chitinase II | 0.02 | 0.03 | 0.02 | 0.02 | 0.01 | 0.02 | 0.69 | 0.01 |
| A6T7X8 | Cob(I)yrinic acid a,c-diamide adenosyltransferase | 0.01 | 0.01 | 0.01 | 0.01 | 0.02 | 0.01 | 1.75 | 0.05 |
| A6T9L3 | Alcohol dehydrogenase | 0.04 | 0.04 | 0.08 | 0.02 | 0.02 | 0.01 | 0.35 | 0.03 |
| A6T9V4 | Glucans biosynthesis protein D | 0.02 | 0.01 | 0.01 | 0.00 | 0.00 | 0.01 | 0.27 | 0.03 |
| A6TA04 | Superoxide dismutase | 0.03 | 0.02 | 0.04 | 0.01 | 0.00 | 0.00 | 0.14 | 0.01 |
| A6TAU7 | Probable formate acetyltransferase 3 | 0.28 | 0.25 | 0.28 | 0.19 | 0.25 | 0.23 | 0.83 | 0.04 |
| A6TCB4 | Phosphoribosylglycinamide formyltransferase | 0.03 | 0.02 | 0.02 | 0.03 | 0.04 | 0.03 | 1.55 | 0.01 |
| A6TCL8 | Signal recognition particle protein | 0.05 | 0.03 | 0.04 | 0.06 | 0.06 | 0.06 | 1.51 | 0.01 |
| A6TD49 | CysJ | 0.03 | 0.02 | 0.02 | 0.00 | 0.00 | 0.00 | <0.01 | <0.005 |
| A6TE01 | Uncharacterized protein | 0.02 | 0.01 | 0.02 | 0.02 | 0.02 | 0.02 | 1.35 | 0.03 |
| A6TEB4 | 3-hydroxy-5-phosphonooxypentane-2,4-dione thiolase | 0.02 | 0.02 | 0.03 | 0.01 | 0.01 | 0.01 | 0.46 | 0.03 |
| A6TEE8 | 2-hydroxy-3-oxopropionate reductase | 0.01 | 0.01 | 0.02 | 0.00 | 0.00 | 0.00 | <0.01 | 0.01 |
| A6TEZ6 | Cyclic AMP receptor protein | 0.45 | 0.32 | 0.26 | 0.47 | 0.47 | 0.54 | 1.44 | 0.03 |
| A6TF07 | CysG2 | 0.01 | 0.01 | 0.01 | 0.01 | 0.01 | 0.01 | 0.73 | 0.02 |
| A6TFY4 | Putative glycoside hydrolase | 0.00 | 0.00 | 0.01 | 0.00 | 0.00 | 0.00 | 0.24 | 0.03 |
| A6TGM2 | NAD(P)H-flavin reductase | 0.06 | 0.05 | 0.05 | 0.08 | 0.08 | 0.08 | 1.54 | 0.01 |
| A6TGU4 | MalE | 0.02 | 0.01 | 0.02 | 0.00 | 0.01 | 0.00 | 0.32 | 0.04 |
| A6TGU6 | LamB2 | 0.30 | 0.28 | 0.41 | 0.18 | 0.22 | 0.26 | 0.67 | 0.04 |
| A6THU9 | 4-hydroxyphenylacetate catabolism | 0.01 | 0.01 | 0.01 | 0.02 | 0.02 | 0.02 | 2.57 | <0.005 |

**Figure 2.**
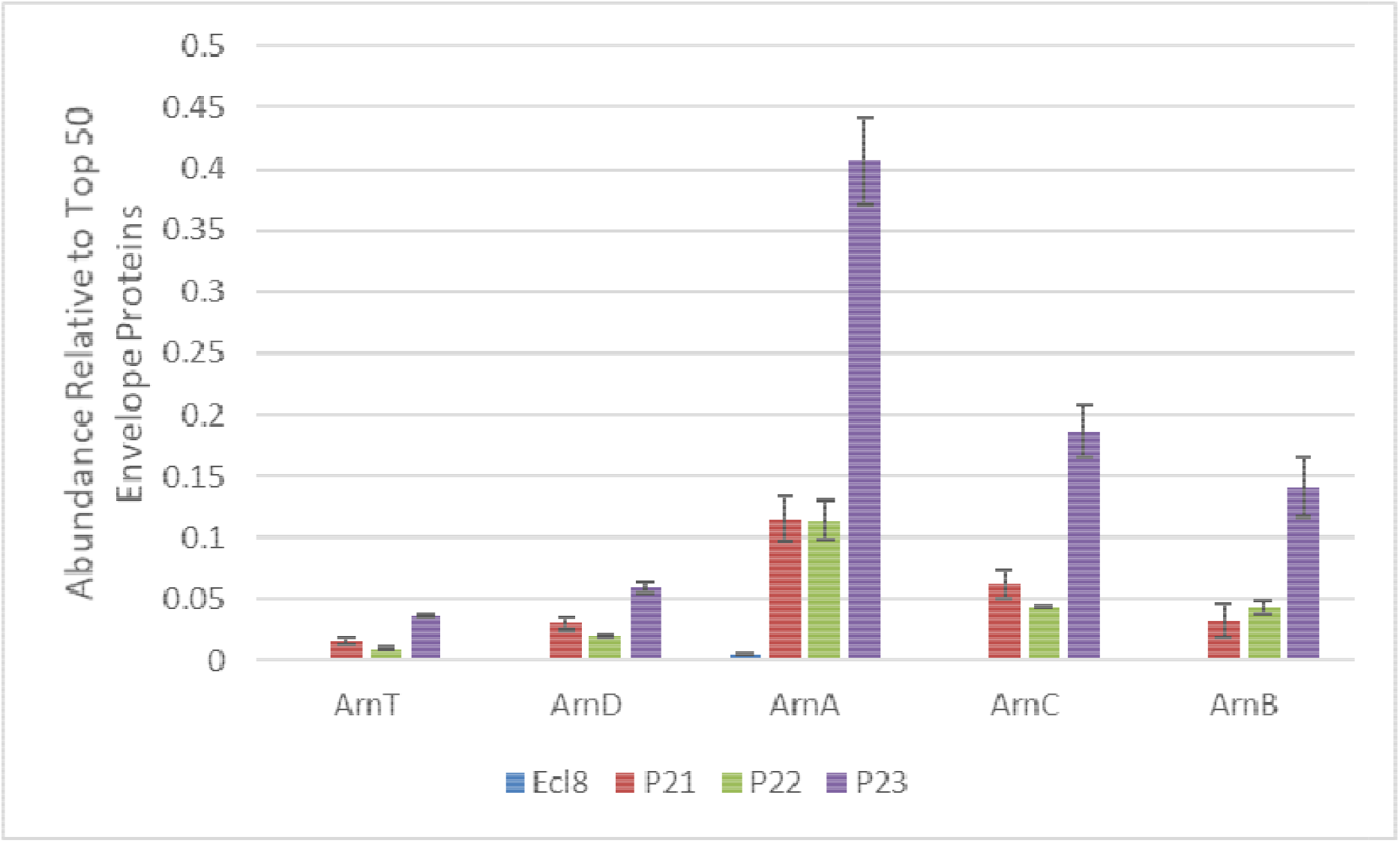
Arn protein abundance in parent strain Ecl8 versus three *mgrB* mutants. Strains were grown in CAMHB and raw envelope protein abundance data for each Arn protein in a sample are presented normalised using the average abundance of the 50 most abundant proteins in that sample. Data for three biological replicates of parent (Ecl8) and *mgrB* mutants (P21, P22 and P23) are presented as mean +/− Standard Error of the Mean. All mutants have statistically significantly increased production of all Arn proteins relative to Ecl8 based on a t-test (p<0.05).

It has been proposed that *arn* operon gene expression is increased upon *mgrB* mutation through activation of the two-component system PhoPQ, and that in addition, there is a secondary increase through activation of the PmrAB two-component system, with PmrD being necessary for transducing PhoPQ activation into PmrAB activation (reviewed in ref. 5). Using *mgrB* mutant P23 as a starting point, we disrupted *phoP*, *pmrD* and *pmrA*, to test the effects of these mutations on Arn protein production and colistin MIC.

Observed Arn protein abundance changes in these regulatory mutants demonstrated the primacy of PhoPQ activation in colistin resistance driven by *mgrB* loss. Arn protein abundance returned to wild-type (**Figure 3**) and colistin MIC fell below the resistance breakpoint (**Table 1**) upon disruption of *phoP* in the *mgrB* mutant P23. In contrast, our proteomics analysis (**Figure 3**) showed that PmrD and PmrAB play only a minor role in increased Arn protein production seen in an *mgrB* mutant. Only ArnC significantly reduced in abundance following disruption of *pmrA*. There was a larger effect following disruption of *pmrD* (4/5 Arn proteins significantly reduced in abundance) but in neither mutant did any of the Arn proteins fall in abundance significantly below levels seen in the *mgrB* mutants P21 and P22 (**Figure 2**) and not surprisingly therefore, the *pmrA* and *pmrD* mutant derivatives of *mgrB* mutant P23 remained colistin resistant, and the MIC against them was only one doubling dilution below that against P23 (**Table 1**). These data add support for our conclusion that there is a threshold of Arn protein abundance required for colistin resistance. It seems clear that PhoPQ activation alone can support an abundance above this threshold even without any additional effects caused by PmrAB activation.

**Figure 3.**
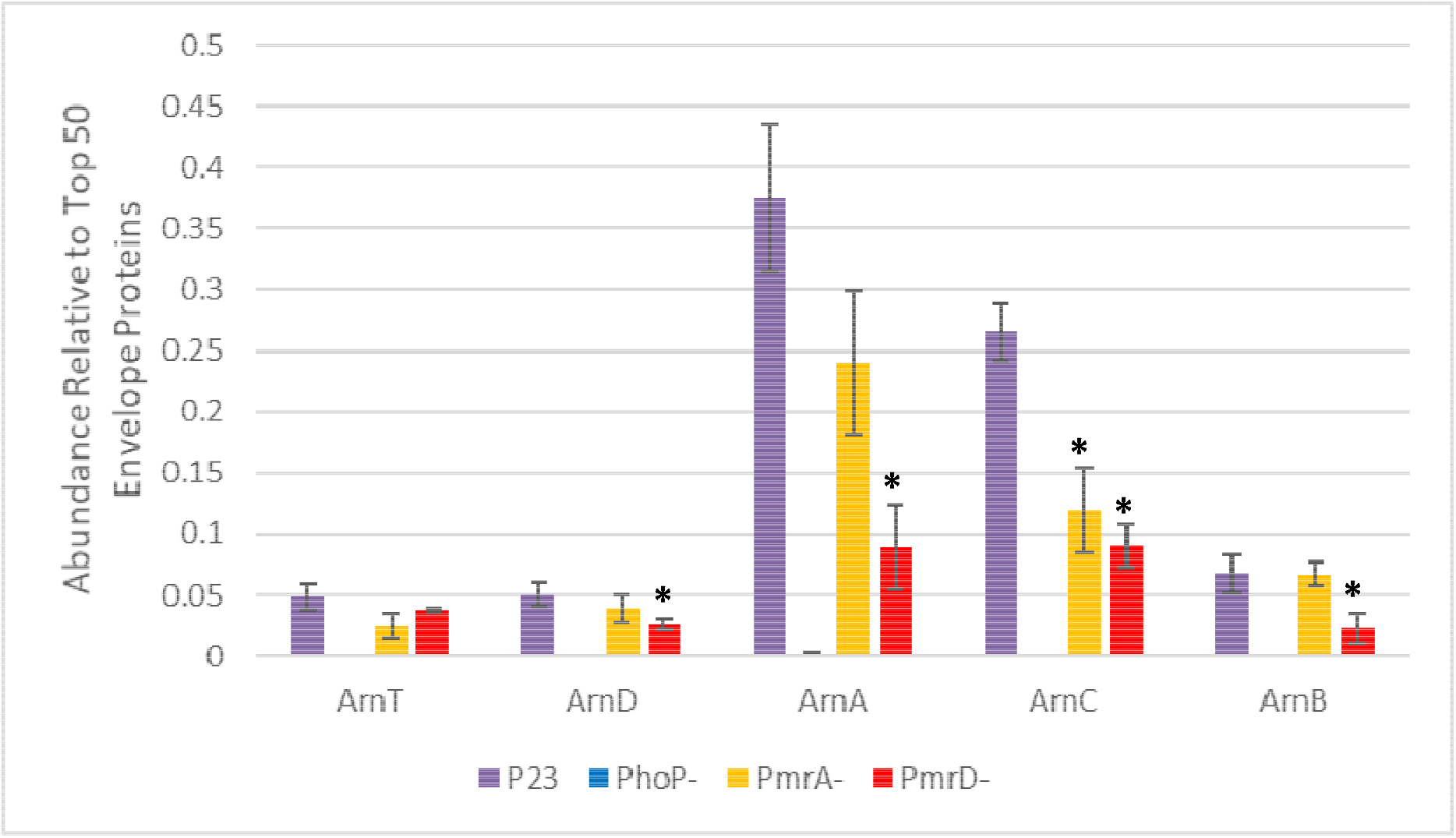
Arn protein abundance in *mgrB* loss-of-function mutant P23 versus its *phoP*. *pmrD* and *pmrA* loss-of-function mutant derivatives. Strains were grown in CAMHB and raw envelope protein abundance data for each Arn protein in a sample are presented normalised using the average abundance of the 50 most abundant proteins in that sample. Data for three biological replicates of parent (P23) and mutants where *phoP*, *pmrD* or *pmrA* had been insertionally inactivated (PhoP-, PmrD- or PmrA-) are presented as mean +/− Standard Error of the Mean. All PhoP-mutants have statistically significantly reductions in production of all Arn proteins relative to P23 based on a t-test (p<0.05). For PmrD- and PmrA-mutants, significant changes relative to P23 are noted with a star.

### PhoPQ regulated proteins identified following mgrB mutation and PhoQ activation

Of 45 proteins (i.e. including the Arn proteins) differentially regulated in all three *mgrB* mutants (**Table 2**) 18 of those upregulated in the *mgrB* mutant P23 returned to wild-type levels upon disruption of *phoP* (**Table 3**). These included the five Arn proteins, the response regulator PhoP itself, LpxO, two Mg^2+^ transporters including MgtA, SlyB and MacA. Transcripts representing all these proteins have been seen to be upregulated in *mgrB* loss-of-function and in PhoQ activatory (PhoQ*) colistin resistant clinical isolates relative to colistin susceptible control isolates through transcriptomics (20). Our proteomics analysis reinforces the definition of a core PhoPQ regulon, but a secondary observation is that the majority (27/45) of the protein abundance changes seen in the *mgrB* mutant P23 do not occur via activation of PhoPQ; they were not reversed following disruption of *phoP* (**Table 3**). The implication of this finding, made here by comparing otherwise isogenic pairs of strains, is that MgrB interacts with regulatory networks other than PhoPQ in *K. pneumoniae*, though these additional effects are not important for colistin resistance, which was completely reversed following disruption of *phoP* (**Table 1**).

**Table 3:** Significant changes in envelope protein abundance seen in *K. pneumoniae* mutant P23 *phoP* versus P23. Strains were grown in CAMHB and raw abundance data for each protein in a sample are presented normalised using the average abundance the 50 most abundant proteins in that sample. Data for three biological replicates of colistin resistant *mgrB* mutant P23 and its *phoP* ertionally inactivated derivative. Proteins listed are those significantly (based on t-test) differently up- (fold change >1) or down-regulated ld change <) in P23 *phoP* versus P23; considering only proteins that were oppositely regulated in P23 compared with its colistin susceptible parent, Ecl8, as listed in **Table 2**. Shading indicates proteins not upregulated in the PhoQ* (activatory) mutant of *K. pneumoniae* clinical isolate 47 relative to KP47; the other 15/18 proteins not shaded are therefore considered the core PhoPQ regulon, see text. Stars indicate proteins coded by transcripts upregulated in a *K. pneumoniae* PhoQ activatory mutant (20).

| Accession | Description | P23 1 | P23 2 | P23 3 | *phoP* 1 | *phoP* 2 | *phoP* 3 | Fold Change | t-test |
| --- | --- | --- | --- | --- | --- | --- | --- | --- | --- |
| A6T4W9 | GlnD | 0.005 | 0.011 | 0.009 | 0.005 | 0.002 | 0.003 | 0.408 | 0.027 |
| A6T5U9 | Putative periplasmic binding protein | 0.036 | 0.029 | 0.024 | 0.000 | 0.000 | 0.000 | <0.01 | 0.001 |
| A6T5Y8 | Mg<sup>2+</sup> transport ATPase* | 0.033 | 0.041 | 0.032 | 0.002 | 0.000 | 0.000 | 0.018 | <0.005 |
| A6T6X6 | MacA* | 0.016 | 0.024 | 0.021 | 0.006 | 0.010 | 0.002 | 0.305 | 0.005 |
| A6T7D1 | Hypothetical Protein | 0.017 | 0.021 | 0.022 | 0.016 | 0.014 | 0.013 | 0.714 | 0.019 |
| A6T7F8 | Thymidylate kinase | 0.021 | 0.018 | 0.016 | 0.016 | 0.013 | 0.011 | 0.740 | 0.039 |
| A6T7J8 | PhoP* | 0.087 | 0.105 | 0.125 | 0.011 | 0.000 | 0.000 | 0.036 | <0.005 |
| A6T9Y9 | SlyB* | 1.833 | 2.191 | 1.513 | 0.669 | 0.634 | 0.697 | 0.361 | 0.002 |
| A6TBQ4 | Lipid A 1-diphosphate synthase | 0.122 | 0.153 | 0.122 | 0.012 | 0.000 | 0.000 | 0.030 | <0.005 |
| A6TBT1 | ApbE* | 0.043 | 0.045 | 0.038 | 0.014 | 0.005 | 0.005 | 0.189 | <0.005 |
| A6TCT2 | LpxO* | 0.048 | 0.052 | 0.037 | 0.009 | 0.000 | 0.003 | 0.081 | 0.001 |
| A6TF96 | ArnT* | 0.040 | 0.070 | 0.035 | 0.000 | 0.000 | 0.000 | <0.01 | 0.006 |
| A6TF97 | ArnD* | 0.071 | 0.041 | 0.041 | 0.000 | 0.000 | 0.000 | <0.01 | 0.004 |
| A6TF98 | ArnA* | 0.496 | 0.309 | 0.321 | 0.000 | 0.000 | 0.000 | <0.01 | <0.005 |
| A6TF99 | ArnC* | 0.236 | 0.313 | 0.247 | 0.000 | 0.000 | 0.000 | <0.01 | <0.005 |
| A6TFA0 | ArnB* | 0.098 | 0.051 | 0.053 | 0.000 | 0.000 | 0.000 | <0.01 | 0.006 |
| A6THH1 | MgtA* | 0.025 | 0.036 | 0.031 | 0.000 | 0.000 | 0.000 | <0.01 | <0.005 |
| A6THT5 | Putative porin | 0.639 | 0.794 | 0.614 | 0.053 | 0.054 | 0.058 | 0.081 | <0.005 |

In order to further investigate the role of direct PhoPQ activation in colistin resistance, we turned to a colistin resistant, PhoQ* (activatory) mutant that we selected from *K. pneumoniae* clinical isolate KP47 (**Table 1**). Whole genome sequencing identified the mutation causes a Tyr89Asn change in PhoQ. Proteomics comparing KP47 with the PhoQ* mutant derivative revealed that levels of Arn protein production in the PhoQ* mutant were not significantly different from those in the *mgrB* loss-of-function mutant P23 (**Figure 4**). Indeed, despite starting with a different parent strain, significant upregulation of 15/18 proteins seen to become downregulated when *phoP* was disrupted in the Ecl8-derived *mgrB* mutant P23 were also upregulated in the PhoQ* mutant relative to its parent, KP47. This further focussed down onto the core *K. pneumoniae* PhoPQ regulon, which is shown in **Table 3**.

**Figure 4.**
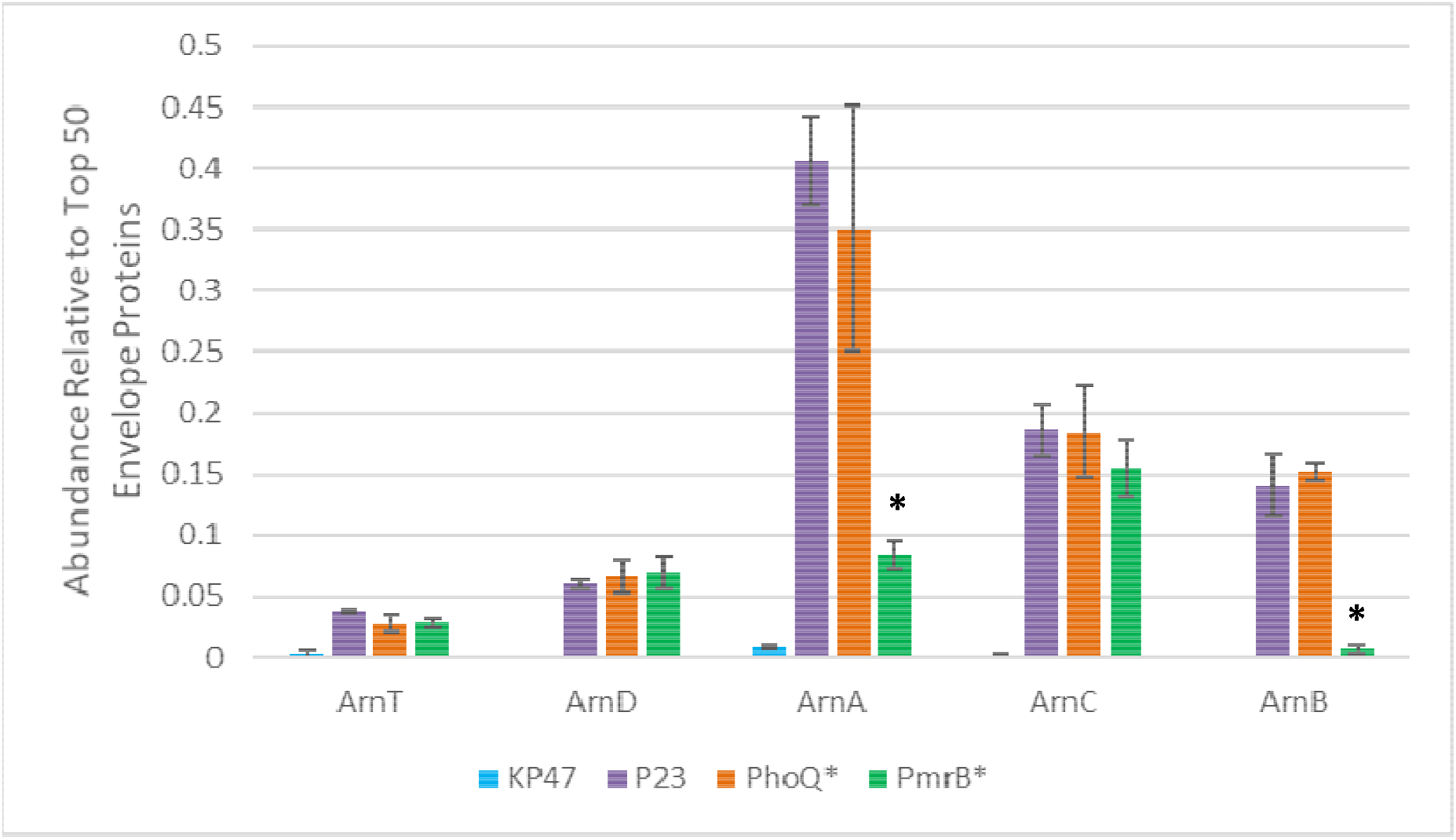
Arn protein abundance in *mgrB* mutant P23 versus clinical isolate KP47 and PhoQ* or PmrB* (activatory) mutant derivatives. Strains were grown in CAMHB and raw envelope protein abundance data for each Arn protein in a sample are presented normalised using the average abundance of the 50 most abundant proteins in that sample. Data for three biological replicates of parent (KP47) and mutants where PhoQ or PmrA have been activated, compared with the *mgrB* loss-of-function mutant P23 are presented as mean +/− Standard Error of the Mean. All Arn proteins, and all Arn proteins except ArnB are significantly upregulated relative to KP47 in its PhoQ* mutant, and PmrB* mutant, respectively based on a t-test (p<0.05). For the PmrB* mutant, significantly lower abundances relative to the PhoQ* mutant are noted with a star.

### Proteins upregulated following activation of PmrAB

Disruption of *pmrA* had little effect on Arn protein production in the *mgrB* mutant P23 (**Figure 3**) suggesting that PmrAB activation is not a major cross-regulatory effect of *mgrB* loss in the context of colistin resistance. Nonetheless, it has been reported that PmrAB activation directly by mutation can confer colistin resistance (23,24) and indeed, we were able to select a colistin resistant mutant of clinical isolate KP47 with an activatory mutation in PmrB (**Table 1**). The mutation identified using whole genome sequencing was Thr157Pro. Arn protein abundance in this PmrB* mutant was significantly elevated relative to KP47 in all cases except for ArnB. For all except ArnA and ArnB the extent of abundance increase was like that seen in the PhoQ* derivative of KP47, and the *mgrB* mutant P23 (**Figure 4**). The fact that ArnB abundance did not increase significantly above the level seen in wild-type KP47 was surprising since the PmrB* mutant is colistin resistant, but it was notable that the MIC of colistin against this PmrB* mutant was one doubling dilution below that against the PhoQ* mutant (**Table 1**). This suggested that either significant upregulation of ArnB is not essential for colistin resistance or there is another mechanism involved in colistin resistance in the PmrB* mutant. In total, only 7/45 proteins significantly up- or down-regulated in the *mgrB* mutant P23 were significantly up- or down-regulated in the PmrB* mutant. As well as the Arn proteins (except ArnB), these were SlyB, LpxO and one Mg^2+^ transporter, which are all part of the core PhoPQ regulon (**Table 2**). Transcripts representing all seven of these PrmAB/PhoPQ dual regulated proteins, plus ArnB have also been seen to be upregulated in clinical isolates with activatory mutations in CrrB (20, **Table 2**) which indirectly activates PmrAB (26,27). In search of an additional mechanism of colistin resistance in the PmrB* mutant we searched the 65 proteins differentially regulated in the PmrB* mutant relative to KP47. Three of those most strongly over-produced were PmrA, PmrB and PmrC (**Figure 5**). Transcription of *pmrC* (also known as *eptA*) is known to be positively controlled by PmrAB in *K. pneumoniae* (24). In *Salmonella* spp. it encodes a phosphoethanolamine transferase, responsible for modifying lipopolysaccharide by decorating it with phosphoethanolamine (25). Importantly, we did not see PmrA, B or C upregulation above our limit of detection (around 100 times less than the level seen in the PmrB* mutant) in any *mgrB* loss-of-function mutant or in the PhoQ* mutant (**Figure 5**). These findings fit, therefore, with our conclusion that cross activation of PmrAB (the direct regulator of *pmrC*) following activation of PhoPQ is very limited under the growth conditions used for our analysis. We also conclude that the PmrAB regulon has limited components that overlap with the PhoPQ regulon, and that PmrC, in terms of functionally important protein product upregulation, is unique to the PmrAB-mediated branch of the colistin resistance-mediating regulatory system. This may well explain why, of multiple studies monitoring the impact of *mgrB* loss-of-function on lipopolysaccharide modification in *K. pneumoniae* (11–13,29) only one has reported elevated levels of phosphoethanolamine modification (29). Indeed, even in this case, contrary to expectations, the observed modification, and the observed upregulation of *pmrC* expression were apparently dependent on PhoPQ but not PmrAB, suggesting it was not caused by PhoPQ-PmrD-PmrAB cross-regulation at all (29). Furthermore, the authors showed that *pmrC* disruption in an *mgrB* loss-of-function mutant background only had a small impact on survival in the presence of colistin (29). This implies that even when rarely seen, phosphoethanolamine modification by PmrC has only a minor role in colistin resistance in *K. pneumoniae*. Indeed, an absence of phosphoethanolamine modification has been advocated as a way to identify mutational colistin resistance (as opposed to *mcr*-mediated resistance, which does cause this modification) whether due to *mgrB* loss-of-function mutation, PhoPQ activation or PmrAB activation in *K. pneumoniae* (30). Overall, this situation is different from that reported in *Salmonella* spp. (25) and another difference between the species is the finding that PmrE production is constitutive in *K. pneumoniae*, and not part of the PhoPQ or PmrAB regulons (**Figure 5**) which is the case in *Salmonella* spp. (5). In the context of PmrB activation in *K. pneumoniae*, this has also been shown by qRT-PCR (26) and in the context of CrrB activation – leading to PmrAB activation – this has been shown through transcriptomics (20). We confirm here using proteomics that PmrE, which is an enzyme responsible for driving the committed step for the biosynthesis of 4‐amino‐4‐deoxy‐L‐arabinose required for colistin resistance (5), is present at levels in wild-type cells similar to the levels of Arn protein produced in colistin resistant mutants (**Figures 4, 5**).

**Figure 5.**
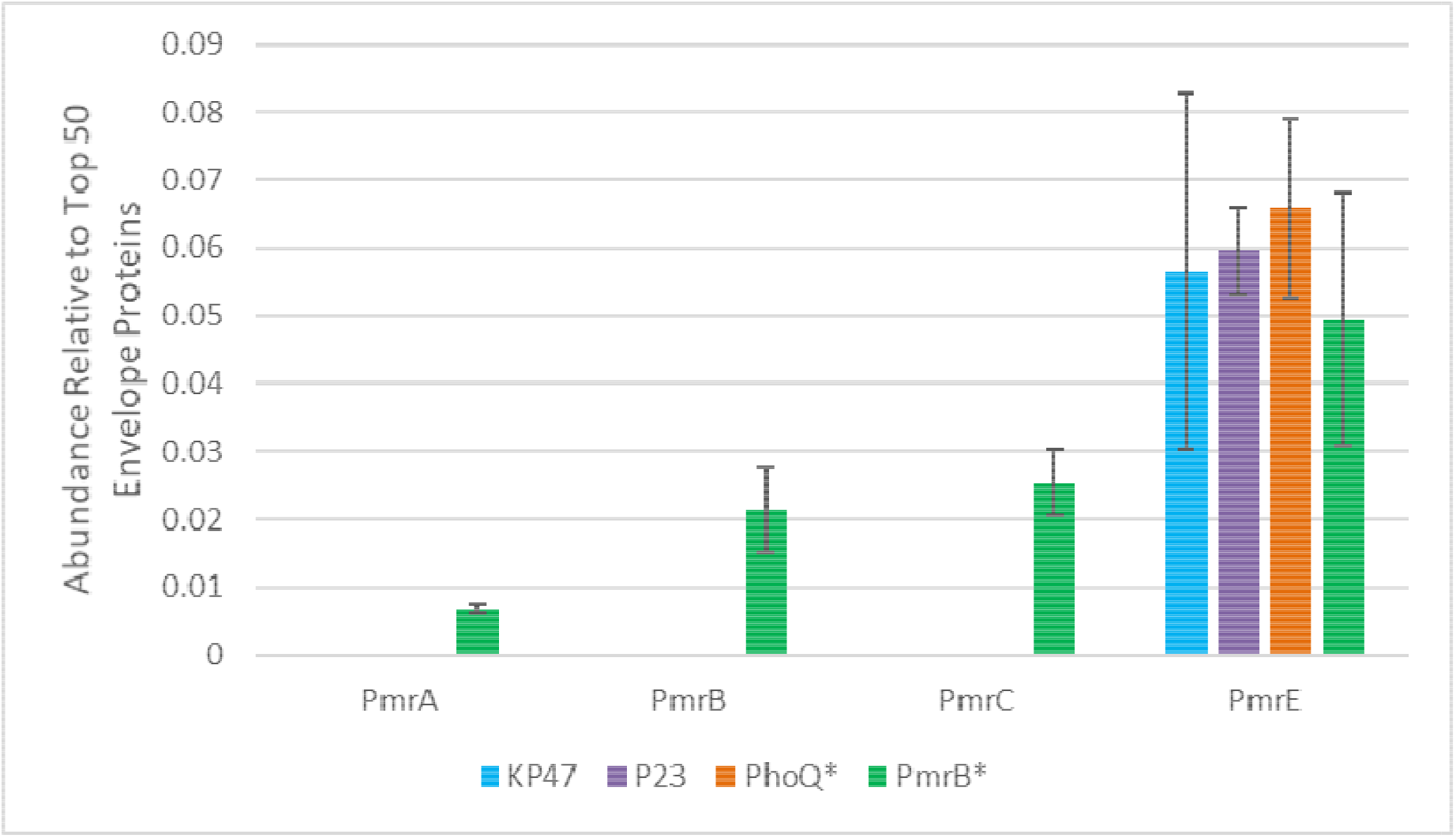
PmrA, B, C and E protein abundance in *mgrB* mutant P23 versus clinical isolate KP47 and PhoQ* or PmrB* (activatory) mutant derivatives. Strains were grown in CAMHB and raw envelope protein abundance data for each Pmr protein in a sample are presented normalised using the average abundance of the 50 most abundant proteins in that sample. Data for three biological replicates of parent (KP47) and mutants where PhoQ or PmrA have been activated, compared with the *mgrB* loss-of-function mutant P23 are presented as mean +/− Standard Error of the Mean. PmrA, B and C were significantly upregulated relative to KP47 in its PmrB* mutant based on a t-test (p<0.05). No other differences were statistically significant.

### Conclusions

Our aim was to definitively investigate the importance of the MgrB-PhoPQ-PmrD-PmrAB signal transduction cascade on Arn and PmrC protein production in *K. pneumoniae*, which has been previously explored using gene knockouts and measurements of transcript levels (5). Based on our data, grounded in protein abundance levels collected during growth in the medium defined for colistin susceptibility testing, we show that this complex interplay, despite remaining potentially important in patients, has minimal importance in the context of colistin resistance in *K. pneumoniae in vitro*. Disruption of *pmrD* or *pmrA* in an *mgrB* loss-of-function mutant does not reduce Arn protein production below a threshold required for colistin resistance (**Figure 3**). Activation of PhoPQ directly or via *mgrB* loss-of-function mutation does not increase the PmrAB-controlled production of PmrC above the level of detection, which is >100-fold less than the observed abundance of PmrC in a PmrB* activatory mutant (**Figure 5**). Therefore, we conclude that colistin resistance caused by PhoPQ activation in conditions defined for colistin susceptibility testing – i.e. in most clinical cases (28) – is due almost exclusively to PhoPQ activation, leading to Arn protein upregulation, and that the level of Arn protein upregulation dictates colistin MIC during growth. It is interesting to note, therefore that clinical isolates with multiple mutations potentially activating PhoPQ can be found, where there is an additive effect on colistin MIC (28). The implication is that real-world colistin usage in the clinical, in some cases, selects for mutations or combinations of mutations that confer colistin MICs above the currently defined resistance breakpoint.

The reason why other earlier seminal reports have placed far higher importance on the cross-activation of PmrAB by PhoPQ via PmrD (14,18) could be that they used media that caused greater basal activation of PmrAB. It is important to remember that the PmrD linker protein has only experimentally been shown to increase the activation of PmrAB once already activated by an external signal, not to activate it from the basal state (14,18). Since PmrAB activation is affected by iron concentration and pH, it may be that the medium used for colistin susceptibility testing – and used by us here – does not activate PmrAB in the first place, so there is nothing that PmrD can do to enhance activation, effectively silencing the cross-regulatory pathway. Another explanation is that previous work relied on measurements of transcript levels. In some cases, small changes in protein abundance are associated with phenotypically relevant changes in antimicrobial susceptibility, as we have shown previously in *K. pneumoniae*, for example upon loss-of-function mutations in *ramR*, which, despite having <5 fold effects on OmpK35 porin and AcrAB-TolC efflux pump production, has large effects on susceptibility to a range of antimicrobial agents from different classes (31). But in some cases, large changes in gene expression are required to have a phenotypic effect when a gene is not highly expressed in the wild-type, for example in the case of OqxAB efflux pump production as controlled by OqxR in *K. pneumoniae*, which needs to increase >10,000 fold to have a phenotypic effect on resistance (32). One major advantage of proteomics is that comparisons of protein abundance can be drawn between different gene products, which is not always the case when transcript levels are measured, since the kinetics of DNA hybridisation can have major influences on signal. This advantage is exemplified here, where we confirmed that *pmrE* expression, measured at the level of protein abundance, is not affected by mutations associated with colistin resistance (**Figure 5**). This was also shown by measuring transcript levels (20,24) but the added value of proteomics is that we found that constitutive PmrE protein levels in all backgrounds are similar to those of the Arn proteins following mutation to colistin resistance, rather than constitutive but at low levels (**Figure 5**).

## Experimental

### Materials, bacterial isolates, selection and generation of mutants

Chemicals were from Sigma and growth media from Oxoid, unless otherwise stated. Strains used were *K. pneumoniae* Ecl8 (30), plus the clinical isolate KP47 (32). To select colistin resistant mutants, one hundred microlitre aliquots of overnight cultures of the parent strain grown in Cation Adjusted Muller-Hinton Broth (CAMHB) were spread onto Mueller-Hinton Agar containing 32 μg.mL^−1^ colistin, which were then incubated for 24 h. Insertional inactivation of *phoP*, *pmrD*, or *pmrA* was performed using the pKNOCK suicide plasmid (33). The *phoP*, *pmrD* and *pmrA* DNA fragments were amplified with Phusion High-Fidelity DNA Polymerase (NEB, UK) from *K. pneumoniae* Ecl8 genomic DNA by using primers *phoP* KO FW (5′-AAGCGGACTACTATCTGGGC-3′) and *phoP* KO RV (5′-TGGAAAGGCTTGGTGACGTA-3′); *pmrD* KO FW (5′-AAGTACAGGACAACGCTTCG-3′) and *pmrD* KO RV (5′-AGTTTATCCCCTTCCCGCAG-3′); *pmrA* KO FW (5’-GACGGGCTGCATTTTCTCTC-3’) and *pmrA* KO RV (5’-TTACCAGGTAGTCATCCGCC-3’). Each PCR product was ligated into the pKNOCK-GM at the SmaI site. The recombinant plasmid was then transferred into *K. pneumoniae* cells by conjugation. Mutants were selected for gentamicin non-susceptibility (5 μg.mL^−1^) and the mutation was confirmed by PCR using primers *phoP* KO RV and BT543 (5’-TGACGCGTCCTCGGTAC-3’); *pmrD* KO FW and BT543; *pmrA* KO RV and BT543.

### Determining minimal inhibitory concentrations (MICs) of colistin

MICs were determined using CLSI broth microtitre assays (34) and interpreted using published breakpoints (35). Briefly, a PBS bacterial suspension was prepared to obtain a stock of OD_600_=0.01. The final volume in each well of a 96-well cell culture plate (Corning Costar) was 200 μL and included 20 μL of the bacterial suspension. Bacterial growth was determined after 20 h of incubation by measuring OD_600_ values using a POLARstar Omega spectrophotometer (BMG Labtech).

### Proteomics

500 μL of an overnight CAMHB culture were transferred to 50 mL CAMHB and cells were grown at 37°C to 0.6 OD_600_. Cells were pelleted by centrifugation (10 min, 4,000 × *g*, 4°C) and resuspended in 30 mL of 30 mM Tris-HCl, pH 8 and broken by sonication using a cycle of 1 s on, 0.5 s off for 3 min at amplitude of 63% using a Sonics Vibracell VC-505TM (Sonics and Materials Inc., Newton, Connecticut, USA). The sonicated samples were centrifuged at 8,000 rpm (Sorval RC5B PLUS using an SS-34 rotor) for 15 min at 4°C to pellet intact cells and large cell debris. For envelope preparations, the supernatant was subjected to centrifugation at 20,000 rpm for 60 min at 4°C using the above rotor to pellet total envelopes. To isolate total envelope proteins, this total envelope pellet was solubilised using 200 μL of 30 mM Tris-HCl pH 8 containing 0.5% (w/v) SDS.

Protein concentrations in all samples were quantified using Biorad Protein Assay Dye Reagent Concentrate according to the manufacturer’s instructions. Proteins (2.5 μg/lane) were separated by SDS-PAGE using 11% acrylamide, 0.5% bis-acrylamide (Biorad) gels and a Biorad Min-Protein Tetracell chamber model 3000X1. Gels were resolved at 175 V until the dye front had moved approximately 1 cm into the separating gel. Proteins in all gels were stained with Instant Blue (Expedeon) for 10 min and de-stained in water.

The 1 cm of gel lane was subjected to in-gel tryptic digestion using a DigestPro automated digestion unit (Intavis Ltd). The resulting peptides from each gel fragment were fractionated separately using an Ultimate 3000 nanoHPLC system in line with an LTQ-Orbitrap Velos mass spectrometer (Thermo Scientific). In brief, peptides in 1% (v/v) formic acid were injected onto an Acclaim PepMap C18 nano-trap column (Thermo Scientific). After washing with 0.5% (v/v) acetonitrile plus 0.1% (v/v) formic acid, peptides were resolved on a 250 mm × 75 μm Acclaim PepMap C18 reverse phase analytical column (Thermo Scientific) over a 150 min organic gradient, using 7 gradient segments (1-6% solvent B over 1 min, 6-15% B over 58 min, 15-32% B over 58 min, 32-40% B over 5 min, 40-90% B over 1 min, held at 90% B for 6 min and then reduced to 1% B over 1 min) with a flow rate of 300 nL/min. Solvent A was 0.1% formic acid and Solvent B was aqueous 80% acetonitrile in 0.1% formic acid. Peptides were ionized by nano-electrospray ionization MS at 2.1 kV using a stainless-steel emitter with an internal diameter of 30 μm (Thermo Scientific) and a capillary temperature of 250°C. Tandem mass spectra were acquired using an LTQ-Orbitrap Velos mass spectrometer controlled by Xcalibur 2.1 software (Thermo Scientific) and operated in data-dependent acquisition mode. The Orbitrap was set to analyse the survey scans at 60,000 resolution (at m/z 400) in the mass range m/z 300 to 2000 and the top twenty multiply charged ions in each duty cycle selected for MS/MS in the LTQ linear ion trap. Charge state filtering, where unassigned precursor ions were not selected for fragmentation, and dynamic exclusion (repeat count, 1; repeat duration, 30 s; exclusion list size, 500) were used. Fragmentation conditions in the LTQ were as follows: normalized collision energy, 40%; activation q, 0.25; activation time 10 ms; and minimum ion selection intensity, 500 counts.

The raw data files were processed and quantified using Proteome Discoverer software v1.4 (Thermo Scientific) and searched against the UniProt *K. pneumoniae* strain ATCC 700721 / MGH 78578 database (5126 protein entries; UniProt accession 272620) using the SEQUEST algorithm. Peptide precursor mass tolerance was set at 10 ppm, and MS/MS tolerance was set at 0.8 Da. Search criteria included carbamidomethylation of cysteine (+57.0214) as a fixed modification and oxidation of methionine (+15.9949) as a variable modification. Searches were performed with full tryptic digestion and a maximum of 1 missed cleavage was allowed. The reverse database search option was enabled, and all peptide data was filtered to satisfy false discovery rate (FDR) of 5 %. Protein abundance measurements were calculated from peptide peak areas using the Top 3 method (36) and proteins with fewer than three peptides identified were excluded. The proteomic analysis was repeated three times for each parent and mutant strain, each using a separate batch of cells. Specific protein abundance was normalised based on the average abundance of the 50 most abundant proteins in each sample. Comparisons of normalised abundance between samples used an unpaired t-test, and significance was defined with p <0.05. Fold-change in abundance between strains was calculated by first calculating average normalised abundance across the three samples representing each strain.

### Whole genome sequencing to identify mutations

Whole genome resequencing was performed by MicrobesNG (Birmingham, UK) on a HiSeq 2500 instrument (Illumina, San Diego, CA, USA). Reads were trimmed using Trimmomatic (37) and assembled into contigs using SPAdes 3.10.1 (http://cab.spbu.ru/software/spades/). Assembled contigs were mapped to the *K. pneumoniae* Ecl8 reference genome (GenBank accession number GCF_000315385.1), obtained from GenBank by using progressive Mauve alignment software (38).

## Acknowledgments

This work was funded by grant MR/S004769/1 to M.B.A. from the Antimicrobial Resistance Cross Council Initiative supported by the seven United Kingdom research councils and the National Institute for Health Research. Genome sequencing was provided by MicrobesNG (http://www.microbesng.uk/), which is supported by the BBSRC (grant number BB/L024209/1).

We declare no conflicts of interest.

## Notes

### Competing Interest Statement

The authors have declared no competing interest.

